# Plant genotype influence the structure of cereal seed fungal microbiome

**DOI:** 10.1101/2022.07.21.500963

**Authors:** Antonino Malacrinò, Ahmed Abdelfattah, Imen Belgacem, Leonardo Schena

**Affiliations:** Dipartimento di Agraria, Università Mediterranea, Reggio Calabria, Italy; Institute of Environmental Biotechnology, Graz University of Technology, Graz, Austria; Leibniz-Institute for Agricultural Engineering Potsdam (ATB) and University of Potsdam, Potsdam, Germany; Agrocampus Ouest, INRAE, Université de Rennes, IGEPP, F-35650 Le Rheu, France

**Keywords:** wheat, ears, cultivar, metabarcoding, ITS, phylosymbiosis

## Abstract

Plant genotype is a crucial factor for the assembly of the plant-associated microbial communities. However, we still know little about the variation of diversity and structure of plant microbiomes across host species and genotypes. Here, we used six species of cereals (*Avena sativa, Hordeum vulgare, Secale cereale, Triticum aestivum, Triticum polonicum*, and *Triticum turgidum*) to test whether the plant fungal microbiome varies across species, whether plant species use different mechanisms for microbiome assembly focusing on the plant ears. Using ITS2 amplicon sequencing, we found that host species influences the diversity and structure of the seed-associated fungal communities. Then, we tested whether plant genotype influences the structure of seed fungal communities across different cultivars of *T. aestivum* (Aristato, Bologna, Rosia, and Vernia) and *T. turgidum* (Capeiti, Cappelli, Mazzancoio, Trinakria, and Timilia). We found that cultivar influences the seed fungal microbiome in both species. We found that in *T. aestivum* the seed fungal microbiota is more influenced by stochastic processes, while in *T. turgidum* selection plays a major role. Collectively, our results contribute in filling the knowledge gap on the wheat seed microbiome assembly and might help in understanding how we can manipulate this process to improve agriculture sustainability.

## Introduction

Plant-associated microbial communities are well known for their impact on the ecology and evolution of their host (Trivedi et al. 2020; Malacrinò, Abdelfattah, et al. 2022). The structure of plant microbiomes largely differentiate within the same plant (e.g., between roots, leaves, fruits, flowers, seeds), and within the same compartment (e.g., different tissues within the same organ) (Trivedi et al. 2020; Dastogeer et al. 2020; Abdelfattah et al. 2016), and it is influenced by several factors, including soil (Zarraonaindia et al. 2015; Malacrinò, Karley, et al. 2021), herbivores (Malacrinò, Karley, et al. 2021; French, Kaplan, and Enders 2021; Malacrinò, Wang, et al. 2021; Frew 2022), pathogens (Ginnan et al. 2020; Wen et al. 2020; Ewing et al. 2021), and abiotic stresses (Vescio et al. 2021; Yu et al. 2022). Despite the great deal of research in this field, we still know little about the rules that govern the assembly of plant microbiomes. Plants, to a certain extent, are able to direct the assembly of their own microbiome. Indeed, plant genotype has been proven to influence, with different strengths, the structure of microbial communities associated with different plant species, including *Boechera stricta* (Wagner et al. 2016), *Medicago trunculata* (Brown et al. 2020), *Glycine max* (F. Liu et al. 2019), *Olea europaea* (Malacrinò, Mosca, et al. 2022), and several others. This effect is thought to occur through changes in the plant metabolome (e.g., exudates, VOCs), and it can be modulated by plants to help coping with biotic and abiotic stresses (H. Liu et al. 2020). However, we are still not able to predict how microbiomes would vary across different plant genotypes.

Most of the research investigating the role of plant genotype on microbiome assembly has been done mostly focusing on roots, leaves, and fruits. However, given the functional importance that microbial vertical transmission can have across generations, it is essential to understand also whether plant genotype influences the composition of the seed microbiome. Few previous studies tested the variation of seed microbiomes across different plant genotypes. For example, Kim et al. (Kim et al. 2020) found that host speciation and domestication shape seed bacterial and fungal communities in rice. Similarly, Wassermann et al. (Wassermann et al. 2022) found different bacterial microbiota associated with different oilseed rape genotypes, with signatures of phylosymbiosis. In addition, while most of the studies constrain their observations to the bacterial and archeal communities (i.e., 16S rRNA gene amplicon metagenomics), it is also essential to focus on the plant-associated fungal communities. Recent research suggests that, within some taxonomical groups, the structure of plant microbiomes reconciliates with the host’s phylogeny (i.e., phylogenetically closer hosts associate with more similar microbial communities). This link, named phylosymbiosis, has been recently found for example in the apples (Abdelfattah, Tack, et al. 2021) and chloridoid grasses (Van Bel et al. 2021). While we still do not know how common is phylosymbiosis among plants, it might be key to understand how different plant genotypes assemble their own microbial communities. This is particularly important considering that the vertical transmission of a portion of the plant microbiome has been found in several species, including oilseed rape (Wassermann et al. 2022), tomato (Bergna et al. 2018), wheat (Walsh et al. 2021), and oak (Abdelfattah, Wisniewski, et al. 2021).

In this study, we tested the influence of plant genotype on the seed fungal microbiome, using cereals as model. We characterized the microbiota of plant ears during the soft-dough phase to best capture the influence of plant genotype on the assembly of seed microbial communities, which might be hindered at a later stage by the plant senescence. First, we focused on the effect of plant species, testing whether the diversity and structure of the fungal microbiome would vary across different cereal species (*Avena sativa, Hordeum vulgare, Secale cereale, Triticum aestivum, Triticum polonicum*, and *Triticum turgidum*). While previous research would suggest driven by plant species, we also expect to detect a signature of phylosymbiosis (the reconciliation of microbiome structure with the host phylogeny). Second, we tested whether different cultivars within the species *T. aestivum* and *T. turgidum* would associate with different fungal communities. According to previous research on wheat (Donn et al. 2015; Azarbad et al. 2020; Yergeau, Quiza, and Tremblay 2020) and other plant species (Wagner et al. 2016; Brown et al. 2020; F. Liu et al. 2019; Malacrinò, Mosca, et al. 2022), we hypothesize to detect a strong genotype-depend signal on the structure of fungal communities within each group.

## Methods

### Field experiment and sampling

For this experiment we selected six plant species: *Avena sativa* (cultivar Argentina), *Hordeum vulgare* (cultivar Pilastro), *Secale cereale* (cultivar Aspromonte), *Triticum polonicum* (cultivar Puglia), *Triticum aestivum* (four cultivars: Aristato, Bologna, Rosia, and Verna), and *Triticum turgidum* (five cultivars: Capeiti, Cappelli, Mazzancoio, Trinakria, and Timilia), with a total of 14 genotypes. All these genotypes were sown during December 2015 in a common garden experiment (each genotype within a 5×5m lot), and ears were harvested in May 2016 during their soft-dough phase. For each genotype, we collected ears from 5 different plants, for a total of 70 samples.

### DNA extraction, library preparation, and sequencing

Samples were lyophilized and grind to a fine powder with stainless steel beads and a bead-beating homogenizer (Retsch GmbH, Haan, Germany). DNA was extracted from ∼40 mg of each sample using the DNeasy Plant Mini Kit (QIAGEN, Venlo, Netherlands) according to the manufacturer’s instructions. DNA quality and concentration was then tested using a Nanodrop 2000 spectrophotometer (Thermo Scientific, Waltham, MA, USA), and samples passing QC were stored at -80°C until further processing.

Libraries were prepared by amplifying the ITS2 region of the fungal rRNA using the primer pair ITS86f and ITS4 (Vancov and Keen 2009). PCRs were conducted by mixing 12.5 *µ*L of the KAPA HiFi Hot Start Ready Mix (KAPA Biosystems, Wilmington, MA) with 0.4 µM of each primer (modified to include illumina adaptors), ∼50 ng of DNA template, and nuclease-free water to a volume of 25 *µ*L.

Reactions were performed by setting the thermocycler (Mastercycler gradient, Eppendorf, Hamburg, Germany) for 3 min at 95°C, followed by 35 cycles of 20 s at 98°C, 15 s at 56°C, 30 s at 72°C, and by a final extension of 1 min at 72°C. A no-template control, in which nuclease-free water replaced the target DNA, was included in all PCR assays. Libraries were checked on agarose gel for successful amplification and purified with an Agencourt AMPure XP kit (Beckman Coulter Inc., Brea, CA, USA) using the supplier’s instructions. A second short-run PCR was performed in order to ligate the Illumina i7 and i5 barcodes (Nextera XT, Illumina, San Diego, CA, USA), and amplicons were purified again as above. Libraries were then quantified using a Qubit spectrophotometer (Thermo Scientific, Waltham, MA, USA), normalized for even concentration using nuclease-free water, pooled together, and sequenced on an Illumina MiSeq (Illumina, San Diego, CA, USA) platform using the MiSeq Reagent Kit v3 300PE chemistry following the supplier’s protocol.

### Data processing and analysis

Paired-end reads were processed using the `DADA2` v1.22 (Callahan et al. 2016) pipeline implemented in R v4.1.2 (R Core Team 2020) to remove low-quality data, identify Amplicon Sequence Variants (ASVs) and remove chimeras. Taxonomy was assigned using UNITE v8.3 database (Nilsson et al. 2019). Reads coming from amplification of plant DNA and singletons were then removed before further analyses.

Data analysis was performed in R v4.1.2 as well. Using the packages `microbiome` (Lahti and Shetty 2019) and `picante` (Kembel et al. 2010) we estimated the diversity of the fungal community for each sample with three different indexes: Faith’s phylogenetic diversity, Shannon’s diversity, and Simpson’s dominance. Then, using the packages `lme4` (Bates et al. 2014) and `car` (Fox et al. 2007), we tested the effect of plant genotype on the three different indexes by fitting three separate linear models specifying *plant species* as fixed factor. The package `emmeans` (Lenth et al. 2019) was used to infer pairwise contrasts (corrected using false discovery rate, FDR).

Similarly, we tested the effect of plant species and cultivar on the structure of seed fungal microbial communities using a multivariate approach with the tools implemented in the `vegan` package (Dixon 2003). Distances between pairs of samples, in terms of community composition, were calculated using a unweighted Unifrac matrix. Tests were run using PERMANOVA (999 permutations), and visualization was performed using a NMDS procedure. First, we tested a model that includes both plant species and cultivar (nested within species) (*∼ plant_species × plant_species/cultivar*). Then, we tested the effect of cultivar within *T. aestivum* and *T. turgidum*, separately. Pairwise contrasts were inferred using the package `RVAideMemoire` (Hervé 2020), correcting p-values for multiple comparisons (FDR).

When testing the influence of plant genotype on the structure of plant microbiota, results suggested that plant species explained a wide portion of the variation (∼ 24.91%, see Results below). To further dissect this result, microbial data were further processed together with an ultrametric phylogenetic tree of Poaceae species obtained from TimeTree (Kumar et al. 2017). A Mantel test (9999 permutations) was used to test the correlation between a Unifrac matrix of the distance between plant species calculated considering the composition of microbial communities (thus, averaged across replicated samples within the same host species) and a matrix of phylogenetic distance between plant species obtained using the function `cophenetic.phylo()` from the `ape R` package (Paradis and Schliep 2019).

Results also showed a greater differentiation within *T. aestivum* compared to *T. turgidum*, suggesting that different mechanisms might contribute to the seed microbiome assembly within each group. We further dissected this by estimating the fungal taxa turnover (taxa replacement) and nestedness (gain/loss of taxa) by partitioning the beta diversity using the package `betapart` (Baselga et al. 2018). Differences between turnover and nestedness were tested, within each plant species, by fitting a linear model. For each cultivar of *T. aestivum* and *T. turgidum*, we tested whether the microbiome assembly fits a null model (Sloan et al. 2006), estimating the goodness of fit to the null model and the immigration coefficient using the package tyRa (https://danielsprockett.github.io/tyRa/).

## Results

Data processing identified 242 fungal ASVs over our 62 samples (Fig. 5), mostly representing the genera *Aureobasidium* (16.19%), *Cladosporium* (15.87%), *Alternaria* (13.54%), *Filobasidium* (12.71%), *Vishniacozyma* (11.82%), *Mycosphaerella* (9.43%), and *Stemphylium* (7.51%).

We first tested whether there is variation in the diversity of fungal microbial communities between the different host plants. We found that host plant species drives an effect on the microbiome diversity, using Faith’s phylogenetic diversity index (F_5, 56_= 7.84, p < 0.001; Fig. 1a), Shannon’s diversity index (F_5, 56_= 9.42, p < 0.001; Fig. 1b), and Simpson’s dominance index (F_5, 56_= 10.57, p < 0.001; Fig. 1c). Post-hoc contrasts clarified that these effects are driven by *A. sativa*, which has a lower diversity (both Faith’s and Shannon’s indexes p < 0.001) and higher dominance (p < 0.001) compared to the other plant species, while no differences were found between the other host plants (p > 0.05).

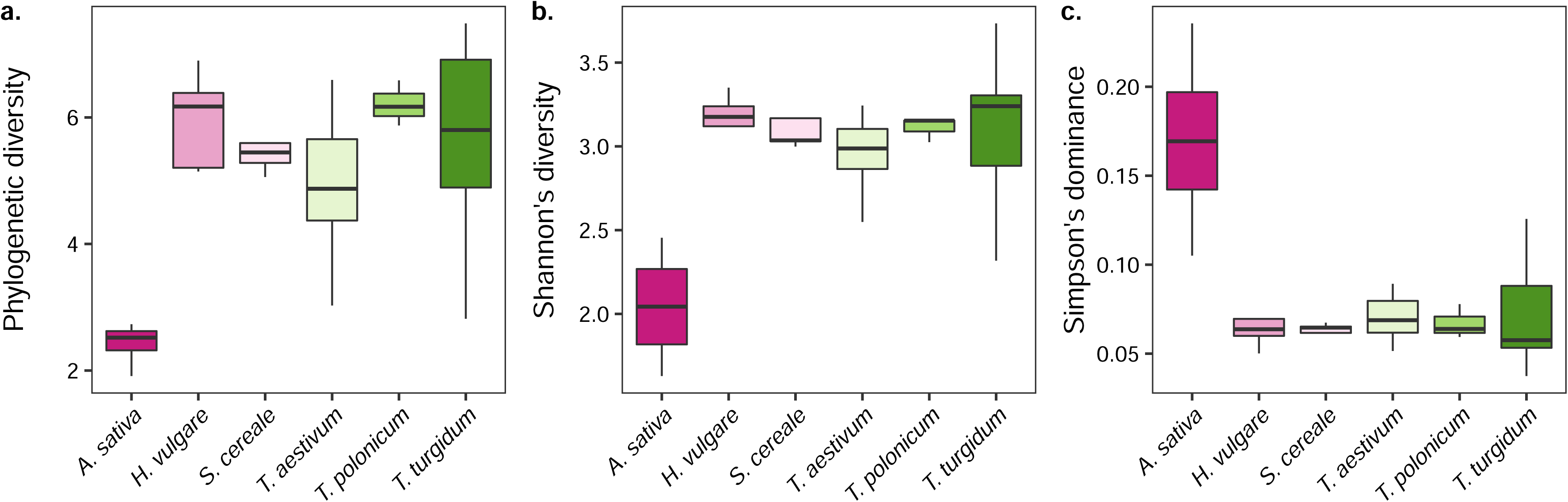

When focusing on the structure of the fungal microbiome, PERMANOVA suggests both an effect driven by host plant species (F_5, 49_= 4.10, p < 0.001) and cultivar (F_7, 49_= 1.83, p < 0.001; nested within species), although species explained a higher proportion of the variance (29.91%) compared to cultivar (15.61%). Post-hoc contrasts show that the structure of the fungal microbiome of *A. sativa* was different compared to all the other plant species (p < 0.05), and the one of *H. vulgare* was different from the microbiome of *T. aestivum*, while no other differences were recorded (Tab. 1).

We then tested whether we could detect a signal of phylosymbiosis, the convergence between host phylogeny and the structure of their microbial communities. Mantel’s correlation shows a weak non-significant correlation between the structure of the fungal microbiota and the host phylogenetic distance (r = 0.31, p = 0.21), although we found overlap across many species when comparing the plant phylogeny with the hierarchical clustering of fungal communities based on Unifrac distances, excluding *H. vulgare* and *T. polonicum* (Fig. 2).

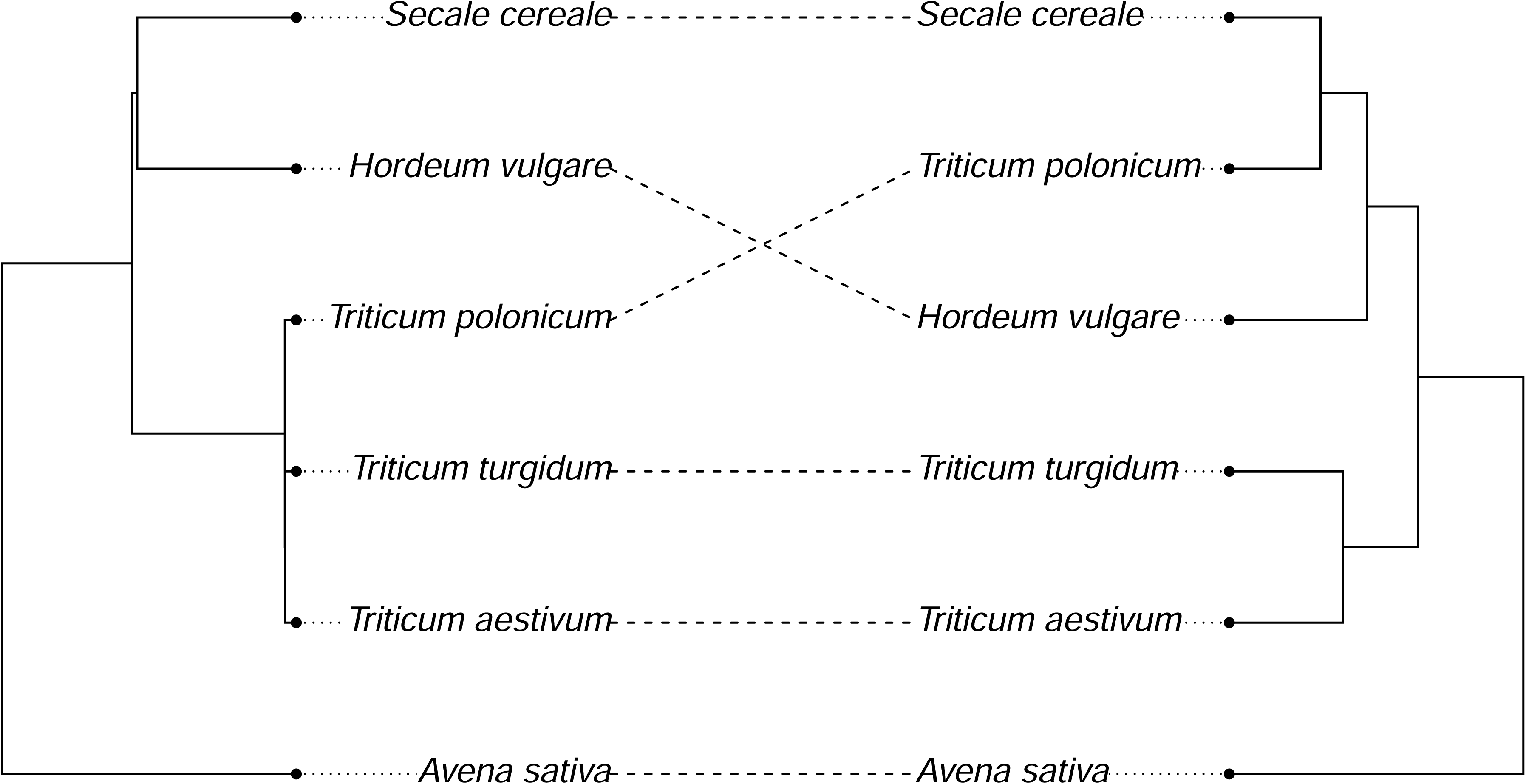

We then tested whether different cultivars within the groups *T. aestivum* and *T. turgidum* associate with a different fungal microbiome. Results suggest an effect driven by cultivar in both *T. aestivum* (F_3, 16_= 1.93, p = 0.005; Fig. 3a) and *T. turgidum* (F_4, 20_= 1.71, p = 0.02; Fig. 3b). Post-hoc contrasts within the *T. aestivum* group show reciprocal differences between the cultivar Bologna, Rosia, and Verna (p = 0.034), while no differences were recorded between the cultivar Aristato and the others (p > 0.05). When testing pairwise differences within the *T. turgidum* group, results did not show any difference between the cultivars (p > 0.05), suggesting that the overall effect driven by cultivar (F_4, 20_= 1.71, p = 0.02) is marginal.

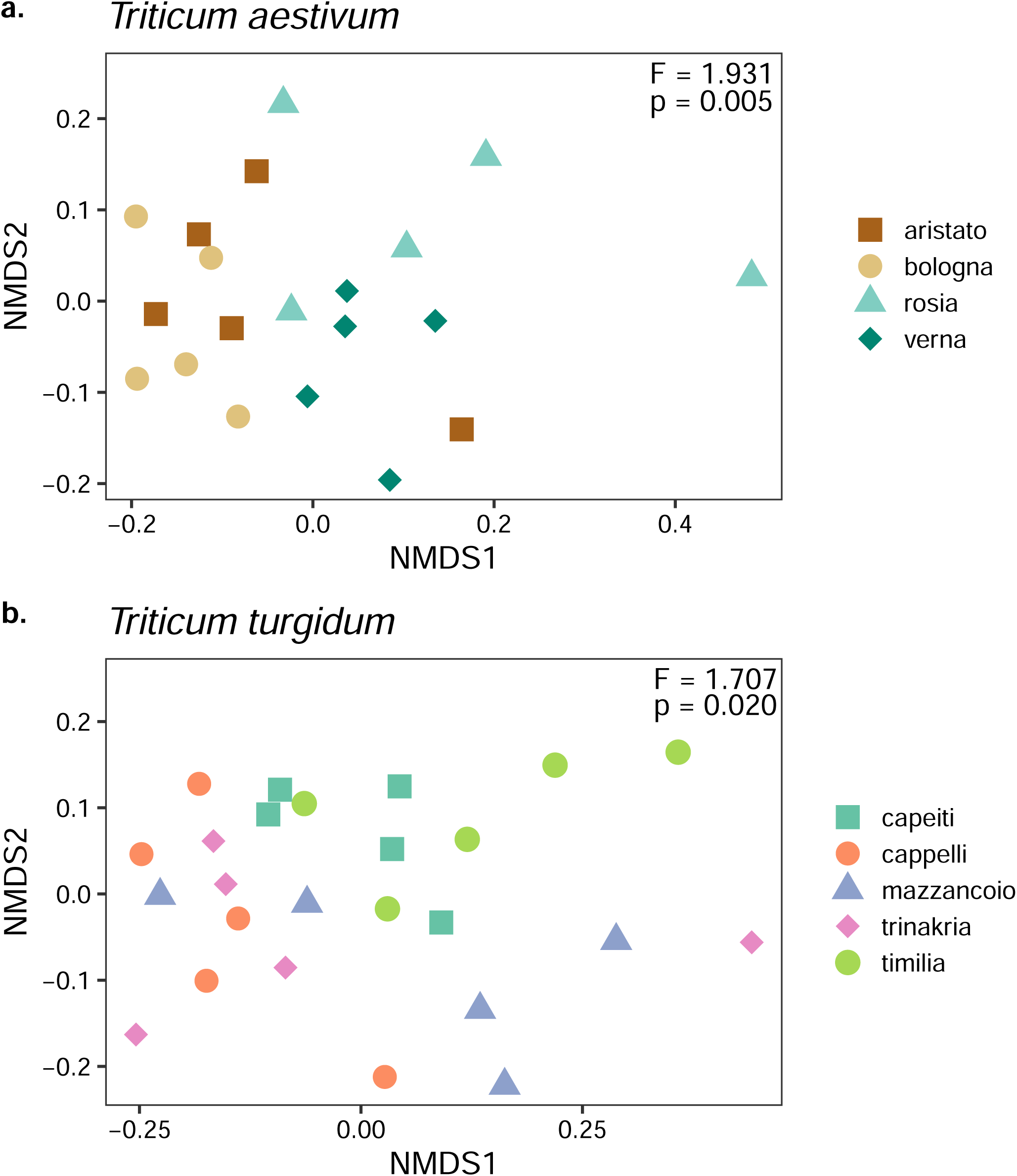

The strongest distinction between cultivars within the group *T. aestivum* compared to *T. turgidum* might suggest that the fungal microbiome of different cultivars assembles by different mechanisms. We tested this idea by estimating the fungal taxa turnover and nestedness within both groups *T. aestivum* and *T. turgidum*. In both *T. aestivum* and *T. turgidum* turnover and nestedness contribute to explain the structure of the fungal microbiome (Fig. 4a). In *T. aestivum*, the contribution of taxa turnover was higher than nestedness (F_1, 378_= 36.31, p < 0.001), while for *T. turgidum* we found the opposite pattern (F_1, 598_= 5.71, p = 0.017). This suggests that in *T. aestivum* there might be a higher replacement of taxa between cultivars, while in *T. turgidum* the fungal community assembles by gain/loss of fungal taxa. This suggests that in *T. aestivum* there might be a higher replacement of taxa between cultivars, probably driven by stochastic processes, while in *T. turgidum* the fungal community assembles by gain/loss of fungal taxa, probably driven by selection. To test for this idea, we fit our data to a neutral model (Sloan et al. 2006), and we estimated the ecological drift (as goodness of fit to the neutral model) and the dispersal (as immigration coefficient from the model).

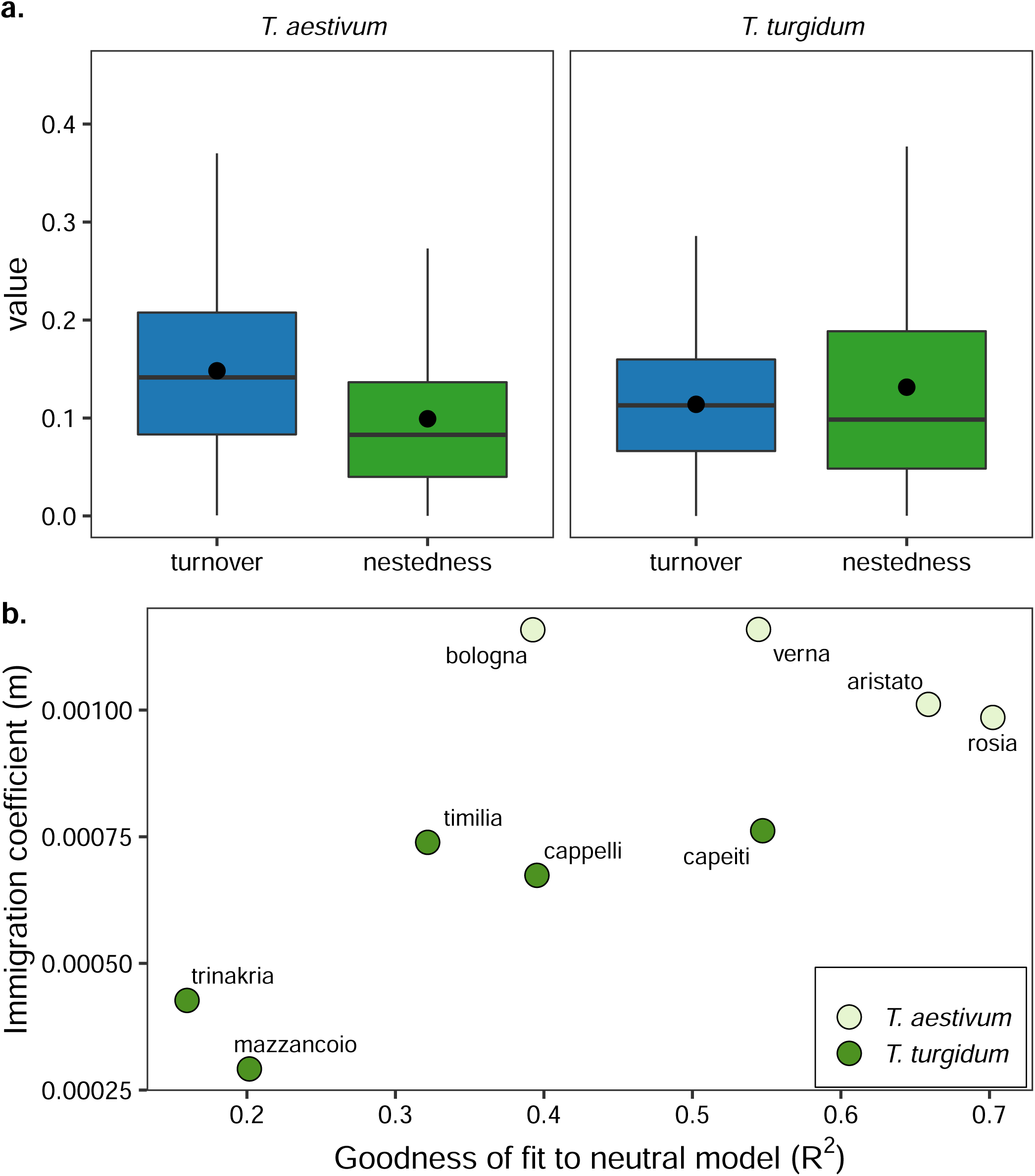

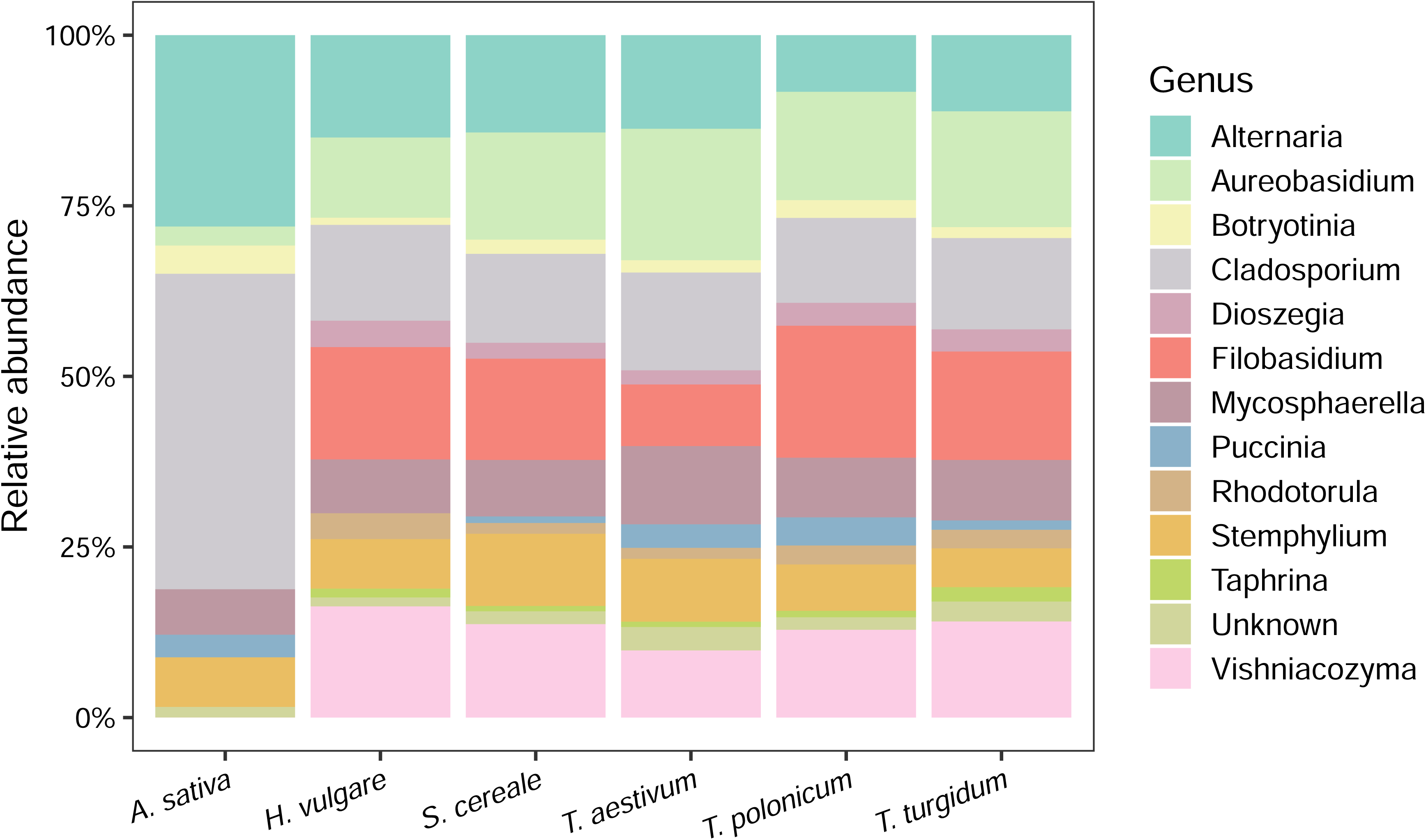

Results (Fig. 4b), show that cultivars of *T. aestivum* tend to have higher R^2^ and dispersal values, while cultivars of *T. turgidum* show the opposite pattern. This suggests that the assembly of seed fungal communities in *T. aestivum* is more driven by neutral processes, while in *T. turgidum* is more driven by selection.

## Discussion

In this study we analyzed the fungal microbial communities associated with ears of six cereal species, and we found variation of the diversity and structure of the microbiome across species, but weak evidence of co-diversification of plant species and microbiota composition. We then focused on the fungal microbiome of different cultivars within the species *T. aestivum* and *T. turgidum*, and we found differences in the microbiota composition between cultivars, with a stronger effect on *T. aestivum* compared to *T. turgidum*. This difference posed the question about the existence of different mechanisms within each plant species when assembling their seed microbiome. Further analyses suggested that in *T. aestivum* the seed fungal microbiome assembly might be driven by the genotype-specific association with fungal taxa, while in *T. turgidum* this assembly might be more driven by stochastic events.

Differences in the diversity and structure of microbial communities between different plant species have been previously reported (Trivedi et al. 2020). In cereals, previous research found evidence of differentiation of the bacterial microbiota between plant species (Kinnunen-Grubb et al. 2020; Wipf and Coleman-Derr 2021; Gholizadeh, Mohammadi, and Salekdeh 2022), while reporting little or no variation in the fungal community (Hassani et al. 2020; Sun, Kosman, and Sharon 2020; Tkacz et al. 2020; Abdullaeva, Ratering, et al. 2021) at different plant compartments. However, to the best of our knowledge, only Abdullaeva et al. (Abdullaeva, Manirajan, et al. 2021) tested this idea on the seed microbiome of different cereal species, suggesting that plant species is a strong driver of the bacterial seed microbiome structure. Similarly, our results show differences in the diversity and structure of fungal microbiome between the different cereal species, although these differences were mostly driven by *A. sativa* and *H. vulgare*. When further digging and testing for the co-diversification of plant species and their fungal microbiome, we found a weak support for phylosymbiosis. This is not surprising, also considering that previous studies focusing on the effect of wheat domestication on the plant microbiome found little or not effect of plant species on the fungal microbiome (Hassani et al. 2020; Sun, Kosman, and Sharon 2020; Tkacz et al. 2020; Abdullaeva, Ratering, et al. 2021), albeit not directly testing for phylosymbiosis.

When further digging into the variation of the seed microbiome between different cultivars, we found that the structure of fungal communities associated with different varieties of *T. aestivum* and *T. turgidum* were shaped by plant genotype. A genotype-driven effect on plant microbiome has been previously shown for several plant species (Wagner et al. 2016; Brown et al. 2020; F. Liu et al. 2019; Malacrinò, Mosca, et al. 2022; Wassermann et al. 2022), but also among wheat varieties (Kavamura et al. 2021). Indeed, previous research found that wheat genotype can shape the diversity and structure of microbial communities in bulk soil (Yergeau, Quiza, and Tremblay 2020), rhizosphere (Donn et al. 2015; Mahoney, Yin, and Hulbert 2017; Azarbad et al. 2020; Kavamura et al. 2020; Rossmann et al. 2020; Simonin et al. 2020; Wolińska, Kuźniar, and Gałązka 2020), roots (Azarbad et al. 2020; Cui et al. 2022), and leaves (Azarbad et al. 2020; Sapkota, Jørgensen, and Nicolaisen 2017; Žiarovská et al. 2020). However, these previous studies mainly focused on *T. aestivum* and on the bacterial communities at each compartment. Latz et al. (Latz et al. 2021) focused instead on the fungal microbiome of seeds, leaves, and roots of 12 cultivars of *T. aestivum*, suggesting that genotype is one of the major driver of the fungal communities. Our results show that such genotype-driven effect occurs in the seed fungal microbiome of *T. aestivum* and *T. turgidum*, with a stronger effect in *T. aestivum*.

These results might suggest different mechanisms in the seed fungal microbiome assembly in *T. aestivum* and *T. turgidum*. When testing this hypothesis, we found that in *T. aestivum* the fungal community has a higher rate of taxa replacement between cultivars (high turnover), a higher fit to a null assembly model, and higher taxa immigration rates, suggesting that neutral processes are the main driver of seed microbiome assembly. On the other hand, in *T. turgidum*, we found a higher rate of taxa gain/loss (nestedness), a lower fit to a null assembly model, and lower taxa immigration rates, suggesting that selection of fungal taxa drives the seed fungal community assembly. This finds support in higher differentiation of fungal communities between cultivars of *T. aestivum* compared to *T. turgidum*, suggesting that in *T. aestivum* the seed microbiome is under the influence of stochastic processes, generating a wider differentiation between cultivars, while in *T. turgidum* selection of the fungal taxa plays a major role, making fungal microbiome assembly more conservative within this group and generating less differences between cultivars. A previous study tested the effect of selection, drift, and dispersal on the assembly of bacterial and fungal communities of rhizosphere, roots, and leaves of *T. aestivum* and its wild ancestors (Hassani et al. 2020), suggesting that in *T. aestivum* the assembly of microbial communities is driven by neutral processes, while in its wild ancestors selection played a major role. Thus, we can speculate that the domestication process somehow influenced the way *T. aestivum* cultivars assemble their microbiome, while this did not happen in *T. turgidum*, and future studies can focus on understanding the reason behind these differences and their functional consequences. For example, the fact that selection of the seed microbiome is more relaxed in *T. aestivum* compared to *T. turgidum* might have consequences on the ability of exterting protection against fungal pathogens (Hassani et al. 2020; Tkacz et al. 2020).

Collectively, our results show that the seed fungal microbiome is variable across cereal species, but also among different genotype of *T. aestivum* and *T. turgidum*, where it is assembled using different mechanisms. While here we are limited on the extent we can infer about microbiome assembly mechanisms and their functional consequences, future studies are needed to finely understand how plants assemble their own microbial communities, how this influence their fitness, ecology, and evolution, and whether we can manipulate the outcome of plant-microbe interactions to improve the agricultural sustainability, ecosystem restoration and natural resources protection efforts.

## Data availability

Raw data is available at NCBI SRA under Bioproject PRJNA848675. The code used to process and analyze the data is available on GitHub.

## Supplementary tables

**Table S1.** Pairwise post-hoc PERMANOVA contrasts between plant species (p-values are FDR-corrected).

| Contrast | p-value |
| --- | --- |
| A. sativa – H. vulgare | 0.0300 |
| A. sativa – S. cereale | 0.0300 |
| A. sativa – T. aestivum | 0.0075 |
| A. sativa – T. polonicum | 0.0600 |
| A. sativa – T. turgidum | 0.0075 |
| H. vulgare – S. cereale | 0.6130 |
| H. vulgare – T. aestivum | 0.0300 |
| H. vulgare – T. polonicum | 0.0575 |
| H. vulgare – T. turgidum | 0.4080 |
| S. cereale – T. aestivum | 0.5862 |
| S. cereale – T. polonicum | 0.4080 |
| S. cereale – T. turgidum | 0.6130 |
| T. aestivum – T. polonicum | 0.4718 |
| T. aestivum – T. turgidum | 0.0862 |
| T. polonicum – T. turgidum | 0.5862 |

